# Dynamic larval dispersal can mediate the response of marine metapopulations to multiple climate change impacts

**DOI:** 10.1101/2020.12.05.413062

**Authors:** Ridouan Bani, Justin Marleau, Marie-Josée Fortin, Rémi M. Daigle, Frédéric Guichard

## Abstract

Climate change is having multiple impacts on marine species characterized by sedentary adult and pelagic larval phases, from increasing adult mortality to changes in larval duration and ocean currents. Recent studies have shown impacts of climate change on species persistence through direct effects on individual survival and development, but few have considered the indirect effects mediated by ocean currents and species traits such as pelagic larval duration. We used a density-dependent and stochastic metapopulation model to predict how changes in adult mortality and dynamic connectivity can affect marine metapopulation stability. We analyzed our model with connectivity data simulated from a biophysical ocean model of the northeast Pacific coast forced under current (1998-2007) and future (2068-2077) climate scenarios in combination with scenarios of increasing adult mortality and decreasing larval duration. Our results predict that changes of ocean currents and larval duration mediated by climate change interact in complex and opposing directions to shape local mortality and metapopulation connectivity with synergistic effects on regional metapopulation stability: while species with short larval duration are most sensitive to temperature-driven reduction in larval duration, the response of species with longer larval duration are mostly mediated by changes in both the mean and variance of larval connectivity driven by ocean currents. Our results emphasize the importance of considering the spatiotemporal structure of connectivity in order to predict how the multiple effects of climate change will impact marine populations.

## Introduction

Climate change has ignited a series of ongoing ecological responses across both terrestrial and marine systems (Parmesan and Yohe 2003, Knapp et al. 2017), and the fastest responses are expected in marine systems (Sorte et al. 2010). Indeed, marine ecosystems are undergoing rapid changes (Doney et al. 2011), including increasing temperature (Walther et al. 2002, Chang et al. 2019), acidification (Hoegh-Guldberg et al. 2007), and rising sea levels (Meehl et al. 2005, IPCC 2013) Species persistence requires local adaptation to changing conditions (Holt and Gomulkiewicz 1997, Hoffmann and Sgro 2011) and/or a range shift (Thomas et al. 2004). For example, many marine species’ thermal tolerances predict that species’ ranges will expand poleward and contract equatorward (Sunday et al. 2012). The range shift hypothesis posits that species may shift their range in response to climate change assuming species can colonize new suitable habitats (Hampe 2004, Guisan and Thuiller 2005, Keith et al. 2008). However, dispersal can be limited by species traits and constrained by physical barriers such as ocean currents (Gaylord and Gaines 2000). Climate change is expected to affect the dispersal process itself (Nathan et al. 2011, Molinos et al. 2017).

The distributions of species can consist of heterogeneous habitats with different levels of suitability and partially connected by dispersal, which can be described as metapopulations of source and sink habitats (Holt 1985, Pulliam 1988, Holt and McPeek 1996, Loreau et al. 2012). In such metapopulations, limited dispersal can promote dynamic stabilization (Andrewarth and Birch 1954, Roff 1974a-b, Bascompte and Solé 1994) and species coexistence (Amarasekare and Nisbet 2001), especially in the presence of spatiotemporal environmental variations (Gonzalez and Holt 2002, Roy et al. 2005).

For many marine species, dispersal occurs during the pelagic larval phase, and regional metapopulation persistence and stability depend on larval transport by ocean currents characterized by strong spatiotemporal heterogeneity (Mitarai et al. 2009, Cowen and Sponaugle 2009; Selkoe et al. 2010). Ocean currents interact with pelagic traits such as pelagic larval duration (PLD) and spawning time (ST) (Watson et al. 2012, Snyder et al. 2014, Bani et al. 2019) to control larval transport and the resulting connectivity among populations. Predicting the response of marine metapopulations to climate change thus requires understanding its direct effects on ocean circulation (Van Gennip et al. 2017) as well as its indirect effect through the plastic response of dispersal traits (Reitzel et al. 2004).

Climate change is expected to affect ocean circulations through changes in oceanic boundary currents (Wu et al. 2012, Hu et al. 2015), which are the main drivers of ocean circulations and by its effects on mesoscale hydrodynamics of tidal residual currents, wind-driven currents, density currents, and gyres (Behera et al. 1999). In general, these changes affect the speed and direction of larvae dispersal and consequently will alter observed patterns of connectivity between marine populations. Climate change also involves increasing temperature, which reduces larval development time and may thus reduce PLD (O’Connor et al. 2007, Munday et al. 2008-2009, Lett et al. 2010). During the adult phase, warming temperature may further impact the reproductive phenology of species by shifting the ST (Kerr et al. 2014, Shanks et al. 2019), reducing reproductive output, and increasing adult mortality (Przeslawski et al. 2008, Munday et al. 2009).

Few studies have looked into multiple climate change effects on pelagic traits and ocean currents. Recent studies have examined temperature-induced changes in ST, PLD and ocean currents (Andrello et al. 2015) and investigated the effects of boundary currents (Coleman et al. 2017) as well as the effects of weather and climatic events (Aguilar et al. 2019), but found little effects on connectivity. Munday et al. (2009) suggested that the shift of PLD may increase self-retention based on the assumption of no change in ocean currents and of increasing chances of settling back to natal habitats with reduced pelagic larval duration. The relationship between PLD and self-retention was supported for species with short PLD (Andrello et al. 2015), but this prediction suggests that increasing self-retention caused by plasticity in PLD under climate change can compensate for climate change-induced adult mortality. Nevertheless, no study has investigated how both local and regional processes interact to affect species stability under climate change and how these effects depend on dispersal strategies.

In this study, we used a stochastic metapopulation model predicting metapopulation stability under spatiotemporal heterogeneity in connectivity (Bani et al. 2019) to study the influence of climate-induced changes in adult mortality, larval duration and dynamic connectivity on regional stability. Our goal is to highlight that multiple statistical components of connectivity (variance, covariance, and realized connections) and their interactions provide key insights needed to understand how ecological processes will respond to ocean currents and temperature stochasticity. We partitioned the interacting effects of adult mortality and of statistical components of connectivity (mean, variance, covariance, and realized connections) on metapopulation stability. Our analysis further assigns these effects to processes mediated by pelagic traits (PLD and ST) or by ocean currents. Finally, we apply our theoretical framework to northeast Pacific coastal habitats using simulated biophysical connectivity under *current* (1998-2007) and projected *future* (2068-2077) ocean circulation scenarios and controlling for PLD and ST.

## Methods

We study how density regulation and dispersal stochasticity affect growth and stability in marine metapopulation model by extending previous models from Bani et al. (2019) to consider effects of climate change on both adult and larval stages. We first study metapopulation stability under interacting effects of key statistical components of fluctuating connectivity (mean, variance, covariance, and realized connections) parameters and adult mortality over a broad parameters space. We then apply the mathematical framework to the northeast Pacific system and study the interacting effects of pelagic traits (PLD and ST), adult mortality and ocean circulation on growth and stability under climate change scenarios where both ocean circulation and temperature are affected. We more specifically study the dual effect of climate on larval dispersal through ocean circulation and through temperature mediated effects on PLD.

Our projected climate change effects on circulations are based on biophysical ocean circulation scenarios along the northeast Pacific over *current* (1998-2007) and projected *future* (2069-2077) periods. The temperature effects are similarly based on scenarios of decreases in PLD (short development time) and increasing adult mortality.

### Metapopulation model

We study the dynamics of a single marine benthic species characterized by adult sedentary and pelagic larval phases and occupying fragmented *s* habitats that are interconnected by time varying dispersal *I*_*ij*_ (*t*) (*i,j* ∈ {1, …, *s*}; the *per capita* immigration rate from habitat *i* to *j* , at time *t*. In each habitat, population dynamics are subject to density-dependence imposed by logistic function with carrying capacity. This assumption accounts for intra-specific competition for resources and negative effects of adults on new recruits.

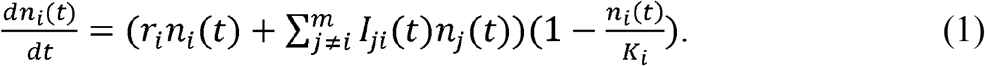

For a population occupying a habitat *i*, *n*_*i*_, represents population density, *K*_*i*_ the carrying capacity, and *r*_*i*_ the local growth rate. Local growth *r*_*i*_ = *Ī*_*ii*_ − *d* results from the balance between adult mortality *d* and time-averaged self-recruitment rate *Ī*_*ii*_ . The time varying per capita immigration rate *I*_*ij*_ (*t*) = *δf C*_*ij*_(*t*) is a function of the proportion *C*_*ij*_ (*t*) of individuals dispersing from *i* to *j*, and of adult fecundity *f* and larval survival rate *δ* , both being uniform over the metapopulation. Both external and local recruitment of larvae are limited by local density, leading to density-dependence acting on self-recruitment and migration rates. The density-dependent model in eq. (1) can be converted to a stochastic multivariate Markovian process (Legault and Melbourne 2019), such that (*x*_l_, … , *x*_*s*_) the vector of relative abundances converges to a stationary distribution given that the deterministic counterpart reaches the stable equilibrium (*k*_1_, … , *k*_*s*_) carrying capacity, where 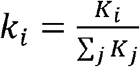. The mean and variance of the stationary distribution of regional relative abundance can then approximated as follow (Bani et al. 2019):

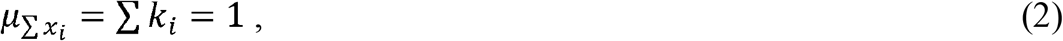

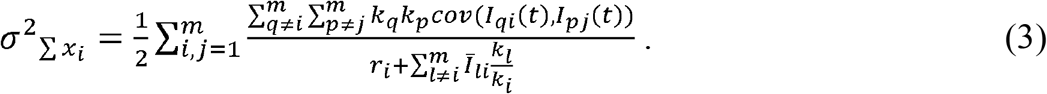

We define stability as the inverse of the coefficient of variation (CV), 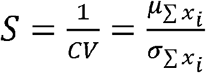, which measures the magnitude of regional relative abundance fluctuation around the total carrying capacity. We used previous analysis of metapopulation stability under dynamic connectivity (Bani et al. 2019) to predict metapopulation response to climate-induced changes in multiple and potential interacting processes including ocean circulation, PLD, and adult mortality. In our case, connectivity from one habitat *i* to another *j* forms a time series (*C*_*ij*_(**t**)={*C*_*ij*_(1),*C*_*ij*_(2), …, *C*_*ij*_(*t*)}) measuring the yearly probability of dispersing. We are then able to calculate the mean connectivity *m* (*μ*(*C*_*ij*_(**t**)), the variance in connectivity *v* (*σ*^2^(*C*_*ij*_(**t**)), the covariance in connectivity between populations *c* (Cov(*C*_*ij*_(**t**),*C*_*kl*_(**t**))) and the number of realized connections between populations relative to the possible number of connections *r*.

### The effects of variations in connectivity and adult mortality

We first use our measure of stability (*S*) to partition the relative importance of the various statistical components of connectivity fluctuations and their interactions with adult mortality. The goal of this analysis is to study how statistical components of connectivity control metapopulation stability through its response to variations in *per capita* immigration (*I*_*ij*_(*t*)) and adult mortality (*d*) . We consider a metapopulation consisting of s = 30 populations and varied adult mortality following a uniform distribution and per-capita immigration rates following a multivariate positive normal distribution while controlling for the mean (vector of size s) and the *s* × *s* covariance matrix. The mean vector and covariance matrix entries are chosen to include the range of simulated connectivity from the biophysical model. We carry out a global sensitivity analysis of stability to connectivity and mortality by calculating the partial ranked coefficient of correlation (PRCC), measuring the association between input variables and their contribution to the output variable. We used the Latin Hypercube Sampling (LHS) method to divide the parameter space into equal probability intervals and compute stability from eq. (3).

### Sources of variations in connectivity

Metapopulation stability *S*(*m,v,c,r,d*) varies with adult mortality *d* and with statistical components of connectivity: the mean m, the variance *v*, the covariance *c* and the realized connections *r*. Stability can thus respond to changes in one or more of these parameters. For instance, we can quantify the stability response to changes in mean connectivity as Δ*S*_*m*_ = *S*(Δ*m*, . , . , . , .) when other statistical components of dynamic connectivity and adult mortality are fixed (see Table 1 for response to other parameters). The net response of stability therefore depends on multiple parameters.

**Table 1:** Components of change in stability and changes in connectivity statistical components associated with abiotic ocean currents and biotic pelagic larval duration (PLD)

|  | Due to<br>abiotic (a)<br>ocean currents change | Due to<br>biotic (b)<br>larval duration reduction |
| --- | --- | --- |
| Stability change due to mean | $\Delta S_{m_a} = S(\Delta m_a, \dots)$ | $\Delta S_{m_b} = S(\Delta m_b, \dots)$ |
| Stability change due to variance | $\Delta S_{v_a} = S(\cdot, \Delta v_a, \dots)$ | $\Delta S_{v_b} = S(\cdot, \Delta v_b, \dots)$ |
| Stability change due to covariance | $\Delta S_{c_a} = S(\cdot, \cdot, \Delta c_a, \dots)$ | $\Delta S_{c_b} = S(\cdot, \cdot, \Delta c_b, \dots)$ |
| Stability change due to realized<br>connections | $\Delta S_{r_a} = S(\cdot, \cdot, \cdot, \Delta r_a, \cdot)$ | $\Delta S_{r_b} = S(\cdot, \cdot, \cdot, \Delta r_b, \cdot)$ |
| Stability change due to adult mortality | $\Delta S_d = S(\cdot, \cdot, \cdot, \cdot, \Delta d)$ | |

Components of dynamic connectivity can change due to changes in ocean circulation (abiotic effect *a*) and in PLD (biotic effect *b*). For example, we can partition changes in mean connectivity into a component Δ*m*_*a*_ driven by changes in ocean circulation and a component Δ*m*_*b*_ driven by changes in PLD (see Table 1 for full list of components). We can use this formalism to study how the complex impacts on multiple components of connectivity and species traits affect our ability to predict potential net effects of climate change on metapopulation stability.

### Pacific northeast and scenarios of ocean circulation

We adopted a biophysical model of larval dispersal for the northeast Pacific to study the effect of pelagic traits and their response to temperature and ocean circulation changes for predicting components of connectivity fluctuations and metapopulation stability under climate change scenarios. We used a solution from a three-dimensional hydrodynamic model ROMS (Regional Ocean Modeling System) output in LTRANS (North et al. 2011) to generate neutrally buoyant particle trajectories along the northeast Pacific coast. The ROMS model uses 3 km horizontal resolution and a 31 vertical resolution levels. The same ROMS model is forced with two different initial conditions. To hind-cast ocean currents over the 10-year period from 1998 to 2007, Masson and Fine (2012) forced the ROMS output with tides, 3-hourly winds from the North American Regional Reanalysis (NARR; Mesinger et al. 2006), heat fluxes, freshwater runoff (Morrison et al. 2011), and lateral boundary salinities and temperature from Simple Ocean Data Assimilation (SODA; Carton and Giese (2008)).

To forecast ocean currents over the 10-year period from 2068 to 2077, Morrison et al. (2014) computed initial and forcing conditions using Canadian Global Climate Model 3 (CGCM3) with the Canadian Regional Climate Model (CRCM). The expected changes derived from this model are an increase in sea surface temperature, increase in sea surface levels, and stronger currents due to fluvial outflows (Foreman et al. 2014). The changes in ocean circulation due to these changes represent our abiotic effect (Table 1).

The two ROMS outputs are used in LTRANS (North et al. 2011) to generate neutrally buoyant particle trajectories and quantify the connectivity (probability of between-patch dispersal). The northeast Pacific coast is divided into 388 (*s* = 388) release sites (a circular site with center on the coastline and radius of 5 km).

During the time period between Jun 04 and Jul 30 of each year from 1998 to 2007, and then from 2068 to 2077, passive and buoyant particles are released at sea floor level biweekly {*Jun* 04, *Jun* 18, *Jul* 02, *Jul* 16, *Jul* 30} from each site, and the locations of each particle is recorded weekly {1,8, … ,113, 120} after release. The release day represents the ST of larvae into water columns while the time spent in water columns before recording the location represent the PLD. For each combination of PLD and ST values, we calculated the time series of all possible pair-wise connectivity probabilities *C*_*ij*_(*t*) of moving from a site *i* to another *j* for all 388 habitats along the northeast Pacific coast (see Bani et al. 2019 for more details).

### Climate change: scenarios of larval duration reduction and adult mortality increase

We assume benthic species with a pelagic larval phase, and with reproduction occurring yearly over a one-day ST window. All larvae reach competency at the PLD. We assume a species is defined by the combination of PLD and ST values and form a homogeneous population in terms of both pelagic traits and its response to increasing temperature, thus ignoring intraspecific variability. This assumption allows capturing the effects of shift in the mean value of traits rather than their distribution or shape.

To study the effect of increasing sea surface temperature, we assume warmer temperature results in reduced PLD (Reitzel et al. 2004, O’Connor et al. 2007). We considered 2 scenarios: (1) each species incurs a 7-day (scenario PLD-7) or (2) a 14-day (scenario PLD-14) reduction of PLD. These scenarios assume a fixed PLD reduction across PLD values, with a 7-day and a 14-day reduction in PLD representing a 25-50% reduction for a species with PLD of 28 days and a 7-14% reduction for a species with PLD of 100 days. This choice follows from constraints of our biophysical model which is set to return the location of each individual larvae weekly after release. Our scenarios are compatible with observed responses of species with relatively short (the sponge *Rhopaloeides odorabile*, Whalan et al. 2008) and long (the plaice *Pleuronectes platessa*, Comerford et al. 2013) PLD with a reported PLD reduction of up to 50% following a 4° to 8° C temperature increase. Additional interspecific variability in PLD response can be expected given that a stronger response is expected from species adapted to colder (e.g. plaice) compare to warmer (e.g. sponge) water (O’Connor et al., 2007). Our scenarios are thus compatible with a minimal increase of 4° C degrees expected for the worst-case climate change scenario (RCP8.5: “business as usual”) by 2100 (Meinshausen et al., 2011).

We used the biophysical connectivity corresponding to each pair of PLD and ST and set adult mortality to *d* = 0.5 to assess changes in regional stability caused by dynamic connectivity following reduction in PLD from each scenario (PLD-7 and PLD-14). We also considered the temperature effect on ST by shifting spanning time two weeks earlier. However, the five spawning times considered in this study had little impact on simulated biophysical connectivity and resulted in little effect on regional stability. To simulate the effects of temperature on increasing adult mortality, we simulated the metapopulation stability using our predicted biophysical connectivity and a 40% increase in *per-capita* adult mortality rate (= 0.7). We report results as the percentage change in stability for each combination of scenarios (PLD-7, PLD-14, mortality, future ocean currents) using regional stability obtained from the *present* connectivity (1998-2007) with adult mortality (*d* = 0.5) as a control.

## Results

### Interacting effects of dynamic connectivity and adult mortality on metapopulation stability

Metapopulation stability with fluctuating connectivity increases with mean connectivity but decreases with increasing variance and covariance of connectivity and with adult mortality (Fig. 1). Adult mortality and statistical components of connectivity interact to affect their net contribution to overall metapopulation stability. In particular, the negative effects of variance (Fig. 1a) and covariance (Fig. 1b) become stronger with increasing mean connectivity and weaker with increasing adult mortality (Fig. 1d). As a result, the effect of mean connectivity becomes weaker with increasing variance (Fig. 1a) and covariance (Fig. 1b) of connectivity.

**Figure 1:**
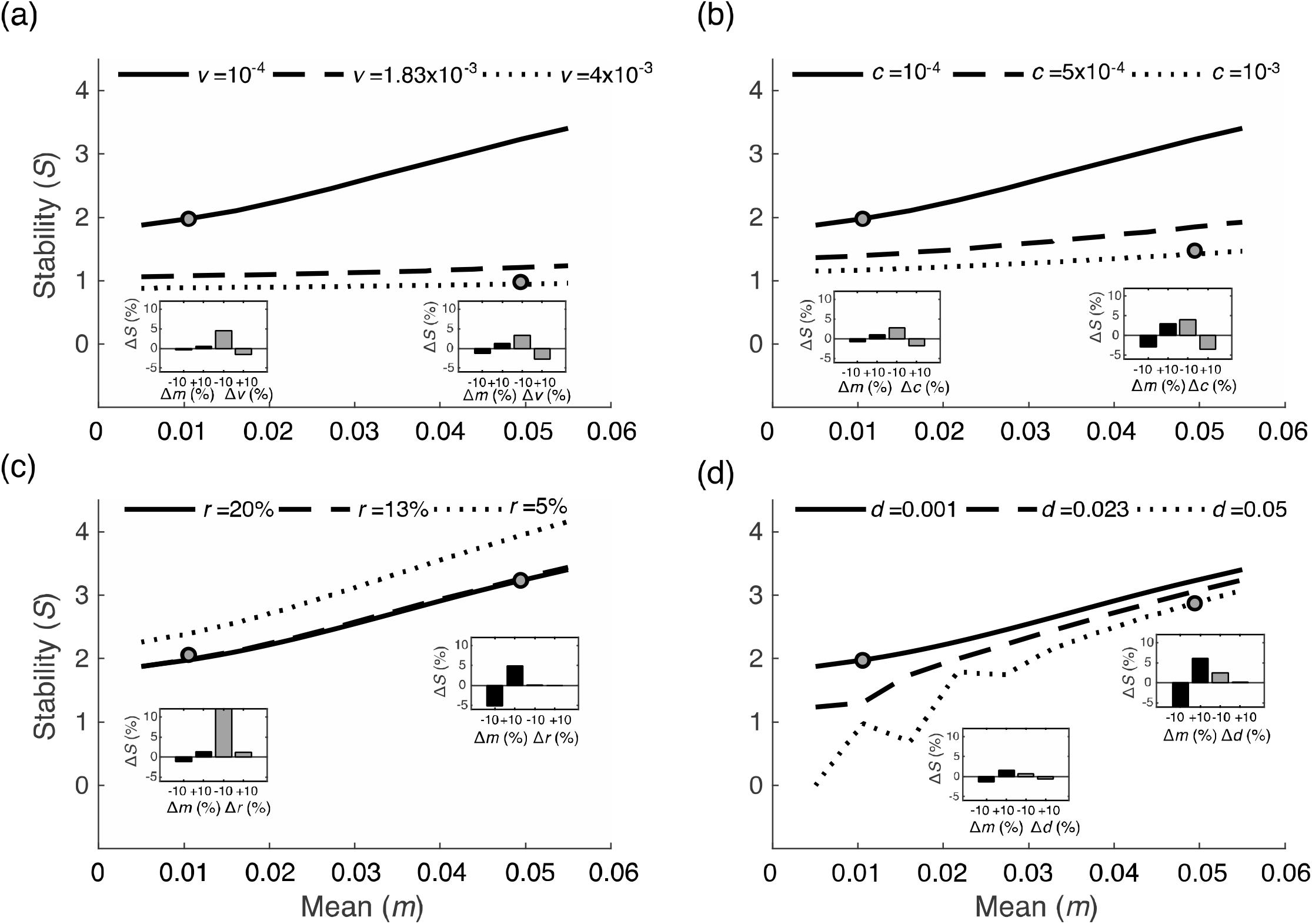
Simulated stability of a metapopulation consisting of *s* = 30 populations as we vary mean connectivity while varying (a) the variance of connectivity, (b) the covariance of connectivity, (c) the realized connections and (d) adult mortality. To highlight the interactions between statistical components of connectivity and adult mortality, in the inner bar plots, the stability changes (Δ*S* in %) with 10 % increase or decrease of connectivity statistical components and adult mortality for two points. One point (left) is chosen at low values mean, variance, covariance and realized connections (*m*=0.0106, *v*=0.00053, *c*= 0.0002, *r*= 6.67%, *d*=0.0064) and the other one (right) is chosen such that mean (m=0.049) and only (a) variance (*v* = 0.0036), (b) covariance (*c*=0.0009), (c) realized connections (*r*= 18.33 %), and (d) adult mortality (*d*=0.0446) are high while the rest components are at same values as the left point.

We illustrate this result by showing how stability can decrease by up to 5% despite a 5-fold increase in mean connectivity when either variance or covariance is also increased (from left to right points, Fig. 1a-d). We can further quantify the response of stability at these points to a ±10% change to each component of connectivity under either low (left points, Fig. 1a-d) or high (right points, Fig. 1a-d) mean connectivity. When mean connectivity is high, changes in variance produce a stronger stability response compared to its response to changes in the mean, realized connections or adult mortality (right inner bar plots in Fig. 1a-d). Moreover, under low mean connectivity, stability increased with both increases and decreases in realized connections (left bar plot in Fig. 1c). This is in contrast with high connectivity cases where positive changes especially in variance increase produce a disproportional decrease in regional stability (right bar plot in Fig. 1a). These results reveal the non-linear effects of multiple statistical components of dynamic connectivity and adult mortality on stability. They show their combined changes can result in contrasting net response of whole metapopulation stability.

### Climate change scenario in the northeast Pacific coastal system

We applied our metapopulation model to the northeast Pacific coast as a case study, both *current* and *future* scenarios, showed that species with short PLD disperse within shorter distances and allow larger mean (*m*) connectivity among the few realized (*r*) connections (Fig. 2). Mean connectivity decreases and realized connections increases with PLD. Moreover, the temporal pattern of larval exchange between connected habitats fluctuates with strong variance (*v*) for short compared with longer PLDs (Fig. 2). The degree of negative (asynchronous) covariance in connectivity between connected habitats quickly decreases with PLD, resulting in random patterns of larval exchange between habitats for PLD > 22 days (Fig. 2). Short PLDs are characterized by a high mean, variance and negative covariance, and all these components are highly sensitive to changes in PLD for PLD < 36 days (Fig. 2).

**Figure 2:**
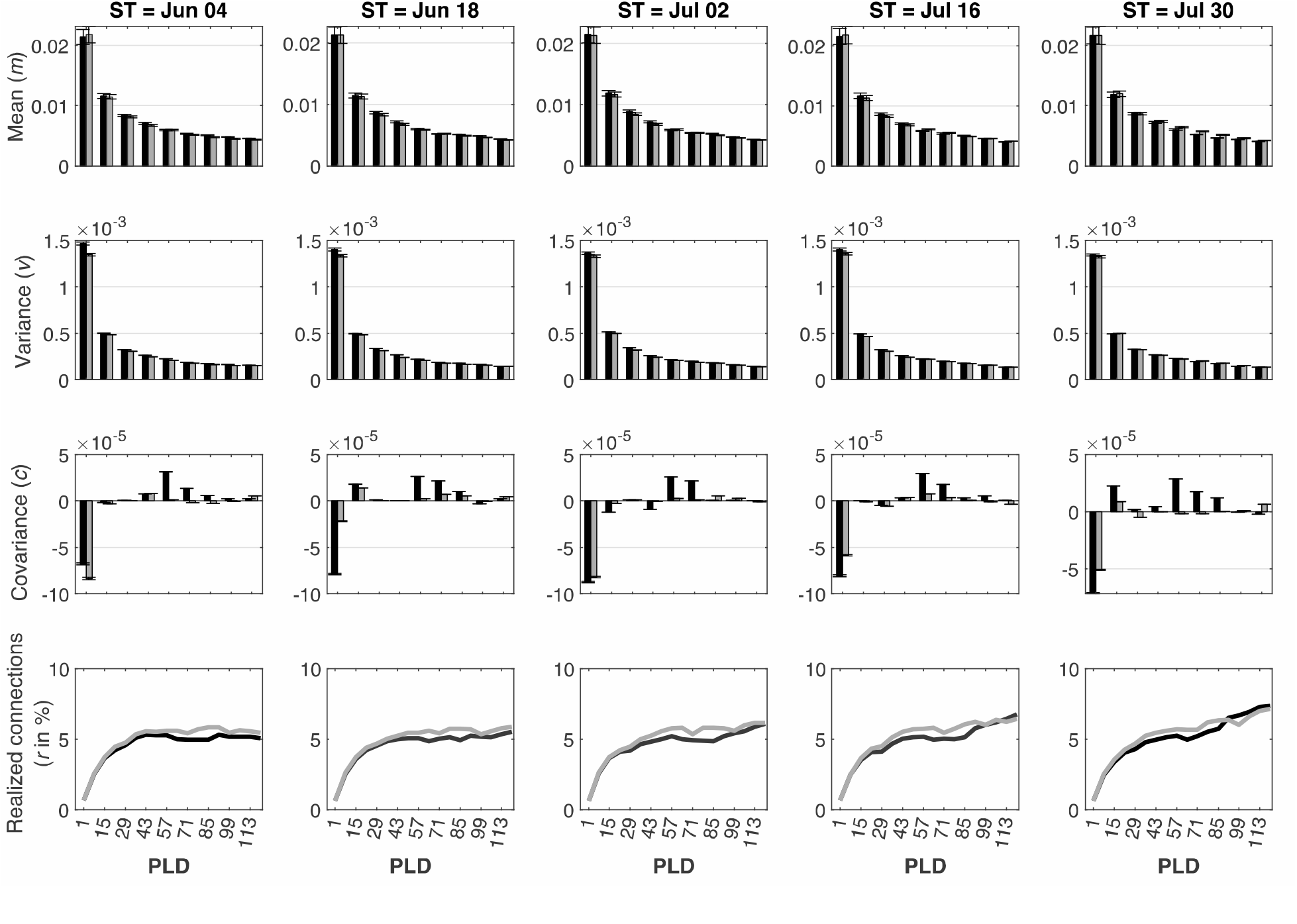
The statistical components of connectivity (rows: mean, variance, covariance, and realized connections) along the BC coastal system and how they vary with PLD (x-axis in unit of days) and the five different STs (columns) for the two ocean currents scenarios: *current* (black) and *future* (light dark). The bars and error bars represent the averages and standard deviations of different connectivity among the 388 habitats along the BC coast.

### Partitioning the effects of multiple statistical components of ocean currents on metapopulation stability

The relationship between PLD and components of connectivity can be further used to partition the relative contribution of individual components to overall metapopulation stability. The mean connectivity and realized connections correlated positively with stability, but the impact of mean connectivity decreases with PLD while the impact of realized connections slightly increases with PLD. The partial ranked correlation coefficient (PRCC) estimating the contribution of mean connectivity to stability decreased with increasing PLD (from 28.78% to 10.37%) while the contribution of realized connections increased from 3.63% to 10.82% (Fig. 3). Both variance and covariance of connectivity correlated negatively with stability and their impact increased with PLD. The negative impact of adult mortality decreased from 61.24% for short PLDs to 42% for long PLDs.

**Figure 3:**
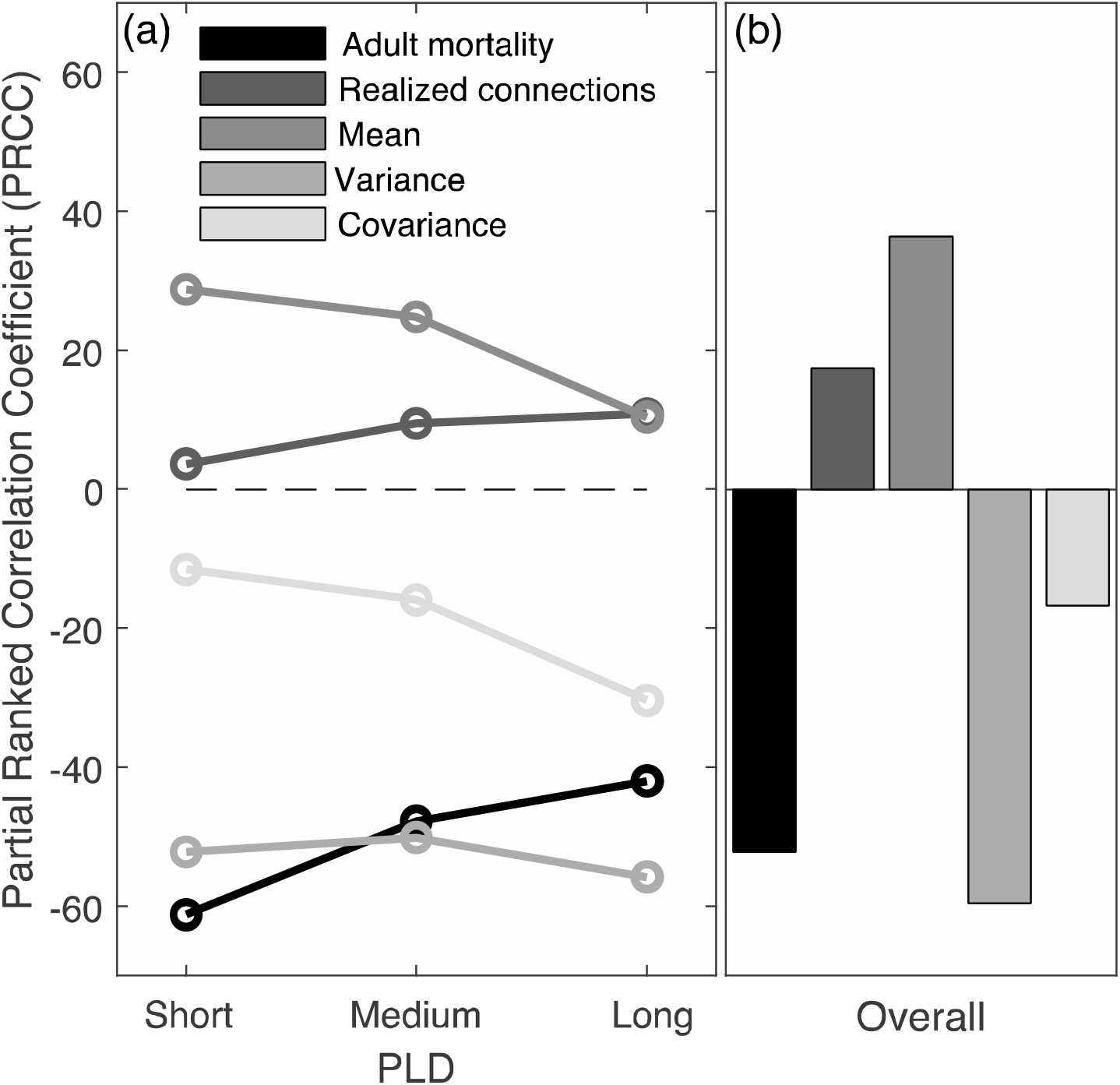
The partial ranked correlation coefficient (PRCC) of statistical components of connectivity and adult mortality based on how it varies depending on (a) the pelagic larval duration (PLD) range group (short:1 to 36 days, medium: 43 to 78 days and long: 85 to 120 days) and (b) over all larval durations.

While adult mortality and variance remain strong drivers of stability across PLD values (Fig. 3b), the strong impact of covariance can only be revealed by resolving its effect in relation to dispersal traits (PLD, Fig. 3). Across our PLD range (1-120 days), regional stability shifts from being more sensitive to mean connectivity, to being more sensitive to covariance (Fig. 3a). This interaction between a biotic trait controlled by temperature, and properties of larval transport that are themselves controlled by climate change, can help predict multiple climate change impacts on metapopulation characterized by dynamic connectivity.

### Effects of changing ocean currents

In general, changes in components of dynamic connectivity between future (2068-2077) and present (1998-2007) scenarios of ocean currents along the northeast Pacific coastal system predict that mean, variance and covariance of connectivity are expected to decrease (Δ*m*_*a*_ < 0, Δ*v*_*a*_ < 0, Δ*c*_*a*_ < 0) and realized connections to increase (Δ*r*_*a*_ > 0) (Fig. 4).

**Figure 4:**
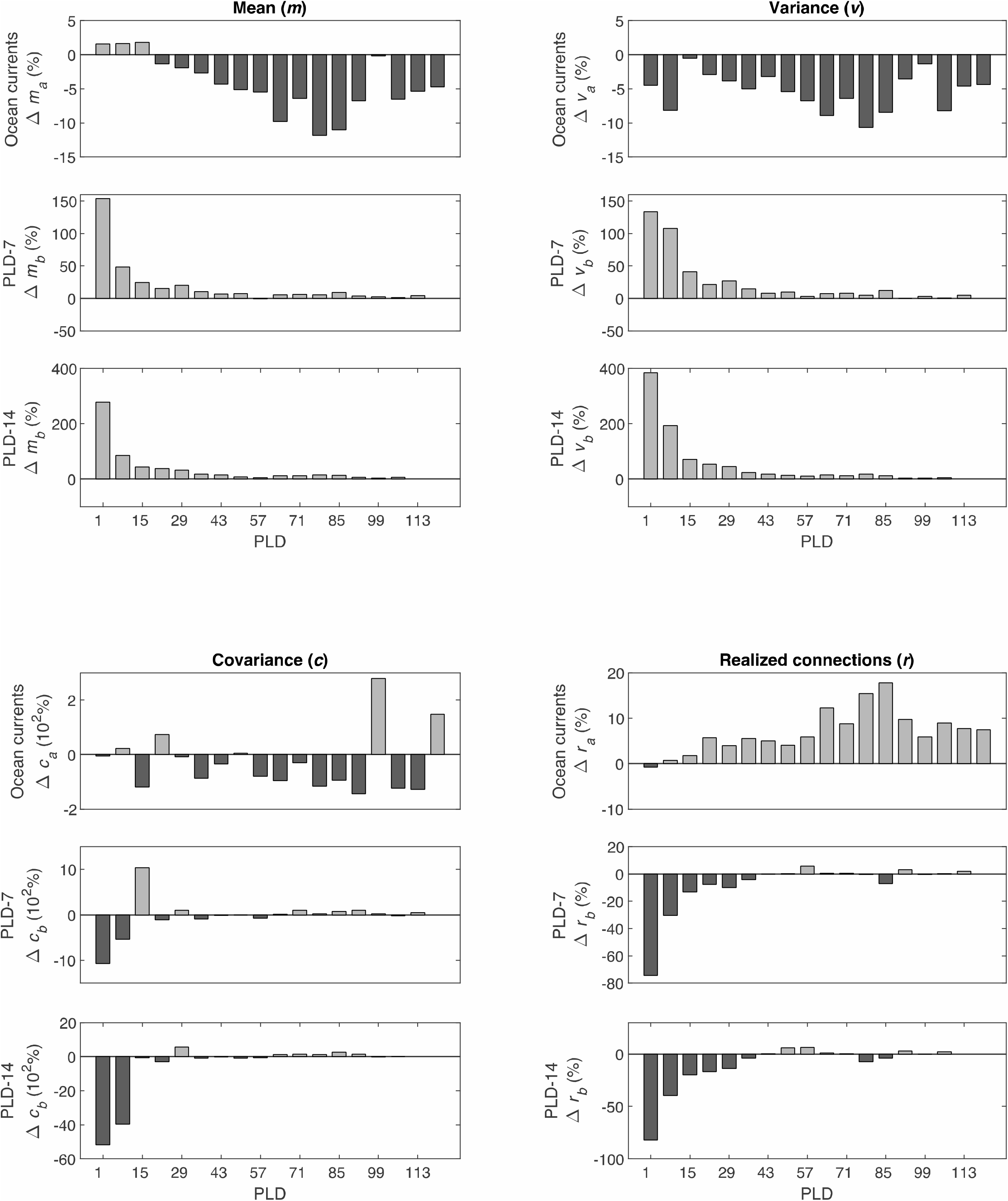
Statistical components of connectivity and how they changed with the different scenarios of abiotic ocean currents and biotic larval duration reduction scenarios PLD-7 and PLD-14. The changes represent the averages among the different STs. Larval duration reduction scenarios (PLD-7 and PLD-14) affect short PLDs and *future* ocean currents scenario affect long PLDs.

Stability increased for the majority of PLDs, especially for longer PLDs (Fig. 5a), which can be explained by corresponding changes in components of connectivity (Fig. 4). The decreases of both mean and variance in our future ocean currents scenario produce a decrease and increase of stability respectively (Fig. 3). These opposite responses to individual components result in a weak predicted change in stability resulting from changes in ocean transport alone (Fig. 5a). However, stability is more sensitive to variance than to mean connectivity (Fig. 3), and the effect of variance decreases still surpassed the decrease in mean connectivity and contributed a net increase of stability across PLD values.

**Figure 5:**
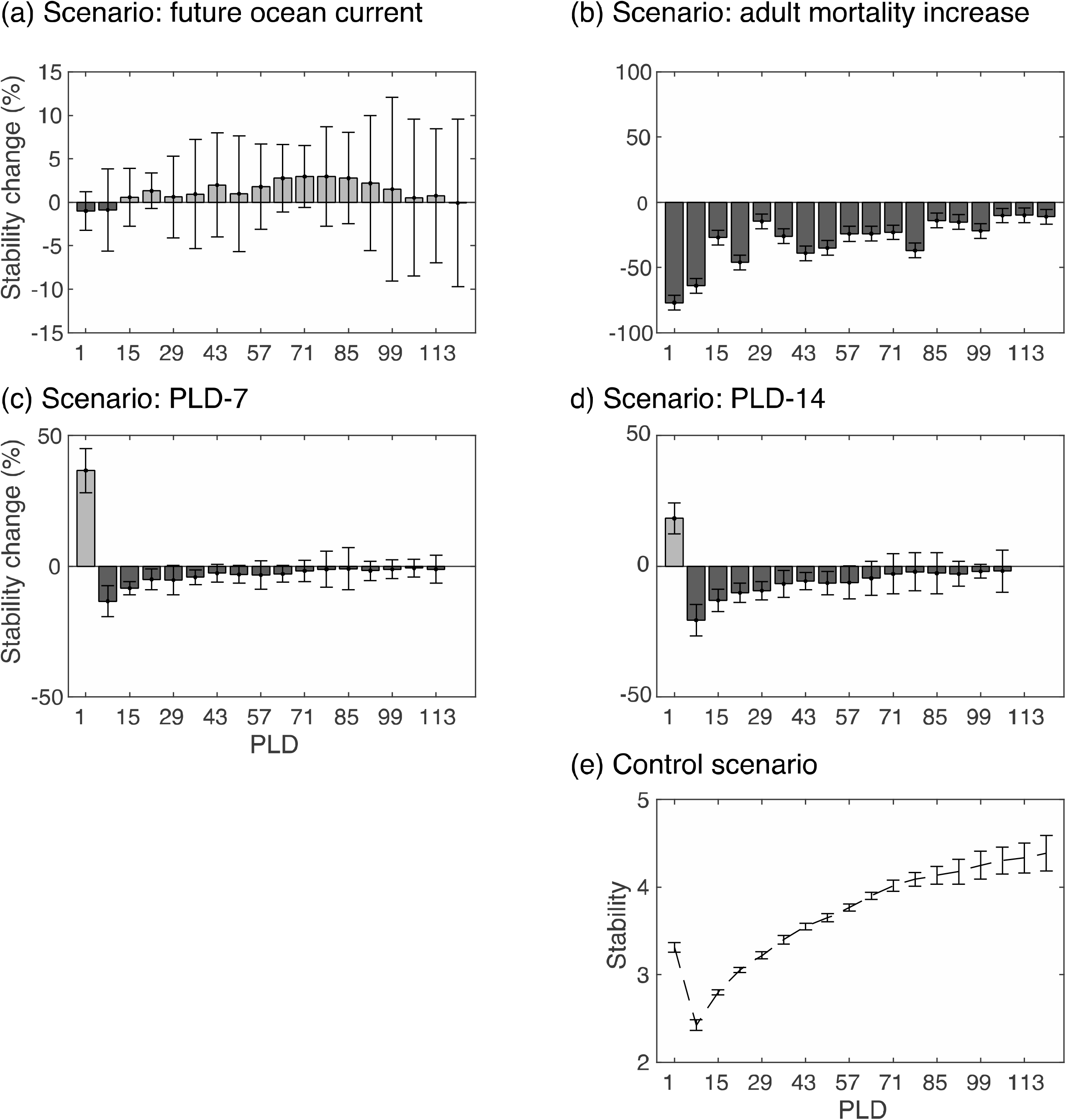
Proportional changes in stability (%) of marine metapopulation under climate change scenarios involving (a) abiotic change in ocean current alone, or the addition of (b) 40% increase of adult mortality, and (c) 7-day and (d) 14-day reduction in PLD. e) Metapopulation stability under the control scenario using present ocean currents biophysical model (1998-2007) and adult mortality at *d*=0.5 is shown as a reference. The bars and error bars show the average and standard deviation among the different five spawning time respectively.

### Effects of changing PLD

The strong effect of PLD on components of connectivity over short PLD values (Fig. 4), directly explain the effect of climate-induced increase in temperature on metapopulation stability (Fig. 5). Both 7-day (scenario PLD-7: Fig. 4) and 14-day (scenario PLD-14: Fig. 4) reductions of larval duration resulted in an increase of mean (Δ*m*_*b*_ > 0t and variance (Δ*v*_*b*_ > 0t of connectivity and in a decrease of realized connections (Δ*r*_*b*_< 0).

In contrast with changes associated with ocean circulation, changes associated with larval duration affect mostly short PLDs. For short PLDs (1-50 days), the reduction of larval duration resulted in an increase of both mean and variance of connectivity (Fig. 4). For PLDs $ 8 days, the temperature driven increase in mean connectivity dominated over both the increase in variance and the decrease of covariance and resulted in an increase of stability in both scenarios PLD-7 and PLD-14 (Fig. 5). For PLDs:2: 8 days, the increase in variance become dominant and resulted in a decrease of stability (Fig. 5).

### Effects of increasing adult mortality

Adult mortality has a negative effect on stability that is opposite to the positive effect of mean connectivity. The sensitivity of stability to adult mortality decreases with PLD similarly to its sensitivity to mean connectivity (Fig. 3). Mean connectivity of species with short larval duration rely on high retention that promotes self-sustainability, but that are prone to longer term local extinction as adult mortality increases. This is in contrast with species with long larval duration which mostly rely on external recruitment and spatio-temporal pattern of dynamic connectivity for regional persistence.

## Discussion

Climate change has multiple interacting effects on metapopulation stability that can be mediated by changes in individual mortality and by spatiotemporal patterns of connectivity among populations. In marine systems, these effects include both direct changes to ocean circulation and individual mortality, and changes in traits such as larval duration that can indirectly affect connectivity by changing how species experience their environment. With simulated larval dispersal from a biophysical model applied to the northeast Pacific forced under current (1998-2007) and future (2068-2077) climate scenarios, along with scenarios of increasing adult mortality and decreasing larval duration induced by temperature, we predicted how species traits (pelagic larval duration) can determine their sensitivity to changes in ocean circulation, and help resolve nonlinear and complex interactions between trait-mediated and environmental processes. Our analysis resolves the relative contribution of these statistical components to regional metapopulation stability across a broad range of dispersal traits values, including pelagic larval duration. These insights are relevant for understanding multiple drivers of climate change impacts on populations across scales. Our results provide a framework for predicting the effects of changes in environmental variability in addition to mean conditions, and changes that affect movement in addition to local individual fitness. They also contribute to current trait-based approaches to climate-change ecology by revealing mechanisms of interactions among multiple traits and environmental drivers.

By considering dynamical properties of connectivity rather than only its mean value, our results can help resolve the effects of individual statistical components of environmental fluctuations that collectively predict metapopulation response. Current metapopulation theories addressing climate-mediated changes in connectivity have largely focused on the physical components of mean connectivity. Studies have, for example, investigated the single-species response to change in temperature mediated by changes in mean connectivity (Andrello et al. 2015, Centina-Heredia et al. 2015, Coleman et al. 2017). However, few studies have addressed trait-mediated responses of connectivity associated with dispersal (Fox et al. 2016, Alvarez-Romero et al. 2018). Our study contributes filling the existing gap in our understanding of how physical and trait-mediated components of marine dispersal can interact to drive metapopulation responses to climate change. For example, our analysis shows that the effects of changes in larval duration on connectivity and metapopulation stability dominate over short larval duration while changes in ocean currents dominate over longer larval duration. Our results thus contribute to a theory integrating physical (transport) and functional (trait-based) metrics of connectivity (Balbar and Metaxas 2019). This integration can be further understood by partitioning the dynamics of connectivity into its statistical components.

The combined and interacting effects of climate change on larval duration and ocean circulation affect the resulting magnitude and direction of changes in each statistical components of connectivity. For example, we found that the temperature-driven reduction of larval duration has effects on dynamic connectivity that are most of the time opposite to the effects of physical transport. Our results emphasize the importance of considering the spatiotemporal structure of connectivity and partitioning this structure into its multiple statistical components. This is important because natural variability in connectivity can be stronger than climate-driven changes in their mean trend (Coleman et al. 2017). Moreover, statistical components of environmental fluctuations such as variance and covariance can be important predictors of metapopulation stability (Bani et al. 2019). Climate-driven changes in PLD and ocean currents translate into strong interactions between the mean, variance and covariance of larval connectivity as they affect metapopulation stability across PLD values.

Our analysis of metapopulation stability over a broad range of PLD values uncovered how its response changes with the changes in the different statistical components of dynamic connectivity as well as with species pelagic larval duration: the stability of species with short pelagic larval duration is more sensitive to the mean value of dispersal among few nearby habitats while species with long pelagic larval duration are more sensitive to temporal variance and covariance of larval dispersal between many distant habitats. Climate-change research in ecology has emphasized the role of connectivity as a mechanism of range shift following increases in temperature (Krosby et al. 2010; Molinos et al. 2017). Our results suggest that changes in mean temperature can affect population persistence through its impact on spatiotemporal connectivity patterns. This result warns against extrapolating and scaling-up individual trait-based response to temperature to whole landscapes (Suarez-Castro et al. 2018).

Our results suggest that dynamic connectivity within metapopulations can also predict population stability and persistence in the face of multiple climate-driven changes in environmental variability. Integrating these roles of connectivity as a driver of local to regional persistence and of larger-scale range shifts will improve of ability to predict shifts in the distribution of species resulting from changes in the mean and (co)variance of the environment. Our results predict long-term response of metapopulation dynamics (stability) to climate scenarios. Integrating dynamic connectivity to our understanding of transient metapopulation response (Holland and Hastings 2008) to ongoing climate change and to its rate (Arumugam et al. 2020) also constitutes an important challenge.

Climate-mediated changes in species traits and environmental variables are key for predicting population response to climate change as they can combine contrasting responses driven by these individual traits and environmental variables. For example, opposite effects of temperature-driven changes in pelagic traits and in physical ocean currents can be observed on larval survival (Centina-Heredia et al. 2015) or spawning time (Andrello et al. 2015). The magnitude and sign of each effect can vary across species and study systems. Our wide range of larval duration values suggests the effects of temperature-driven changes in larval duration should be important for short larval duration compared with physical ocean currents that can affect species with longer larval duration.

This prediction is in agreement with Kendall et al. (2016) who reported that PLD reduction to 10 days explained most of the mean connectivity changes while physical transport explained changes in connectivity associated with longer PLDs (50 and 100 days). This prediction is also compatible with the observed response of specific taxa in relation to their life-history: the smaller effects of physical transport compared to dispersal traits for the dusky grouper, a fish with a short larval duration (Andrello et al. 2015), and the large effects of physical transport for species with long larval duration such as lobsters (6-12 months; Centina-Heredia et al. 2015) and sea urchins (Coleman et al. 2017). While these results apply mostly to the response of mean connectivity, our results also contribute to resolving population responses driven by climate-mediated changes in spatiotemporal patterns of connectivity.

By applying our metapopulation theory to the northeast Pacific coast, we showed how changes of ocean currents and larval duration mediated by climate change interact in complex and opposing directions to shape local mortality and connectivity with nonlinear effects on regional metapopulation stability. Our study contributes to an extension of metapopulation theories to fluctuating and spatially-structured connectivity, and our qualitative predictions based on our case study illustrates the important role of dynamic connectivity for climate change research. However, our more specific predictions are expected to be strongly dependent on characteristics of our study system and on assumptions of our models. Ocean currents are of course highly variables across regions and their complexity is not fully captured by available ocean circulation models. Latitude and the range of temperatures experienced by populations can also have a strong impact on projected temperature increases, and on the response of PLD (O’Connor et al. 2007), which would affect the relative impact of PLD on changes in dynamic connectivity.

Climate change represents a challenging issue for marine protected area (MPA) planning (McLeod et al. 2009, Magris et al. 2014), and recent studies have called for dynamic MPA strategies that can provide and allow species to colonize suitable protected habitats under changing climate (D’Aloia et al. 2019). Based on our findings, mitigating climate change impacts by considering changes in local habitats may apply to only some species. Instead, the design of MPA networks could have to consider the biotic and abiotic contexts leading to synergistic or antagonistic effects of ocean currents and of species pelagic traits on metapopulation stability.

More generally, our ability to mitigate climate change impacts in variable and spatially structured environments might depend on conservation strategies that integrate dynamic connectivity with multiple climate-driven changes in of local habitats, species traits, and environmental heterogeneity.

